# The effect of recombination on the evolution of a population of *Neisseria meningitidis*

**DOI:** 10.1101/2020.04.08.031906

**Authors:** Neil MacAlasdair, Maiju Pesonen, Ola Brynildsrud, Vegard Eldholm, Paul A. Kristiansen, Jukka Corander, Dominique A. Caugant, Stephen D. Bentley

## Abstract

*Neisseria meningitidis* (the meningococcus) is a major human pathogen with a history of high invasive disease burden, particularly in sub-Saharan Africa. Our current understanding of the evolution of meningococcal genomes is limited by the rarity of large-scale genomic population studies and lack of in-depth investigation of the genomic events associated with routine pathogen transmission. Here we fill this knowledge gap by a detailed analysis of 2,839 meningococcal genomes obtained through a carriage study of over 50,000 samples collected systematically in Burkina Faso, West Africa, before, during, and after the serogroup A vaccine rollout, 2009-2012. Our findings indicate that the meningococcal genome is highly dynamic, with recombination hotspots and frequent gene sharing across deeply separated lineages in a structured population. Furthermore, our findings illustrate the profound effect of population structure on genome flexibility, with some lineages in Burkina Faso being orders of magnitude more recombinant than others. We also examine the effect of selection on the population, in particular how it is correlated with recombination. We find that recombination principally acts to prevent the accumulation of deleterious mutations, although we do also find an example of recombination acting to speed the adaptation of a gene. In general, we show the importance of recombination in the evolution of a geographically expansive population with deep population structure in a short timescale. This has important consequences for our ability to both foresee the outcomes of vaccination programmes and, using surveillance data, predict when lineages of the meningococcus are likely to become a public health concern.

## Introduction

*Neisseria meningitidis*, the meningococcus, is a species of bacteria found exclusively in humans. It can cause meningitis: an infection of the membranes covering the brain and spinal cord, as well as septicemia (Stephens et al., 2007). These infections are difficult to treat, even with antimicrobials, and have a high case fatality rate. Of the 12 serogroups defined on the basis of the structure of the capsular polysaccharide, six (A, B, C, W, X, and Y) are responsible for nearly all cases of invasive meningococcal disease (IMD) world-wide. In contrast to strains that are capable of causing disease, non-disease causing carriage isolates are typically unencapsulated. However, most infections, of encapsulated and unencapsulated *N. meningitidis*, are asymptomatic, with the bacteria being carried in the oropharynx of human populations without causing disease with a prevalence of approximately 5-10% (Christensen et al., 2010). It is likely that essentially all individuals will be colonised by potentially IMD-causing bacteria once or even several times during their lifetimes, so there are an uncertain number of carriage infections and transmission events in a human population. This presents a challenge for controlling the disease, and in order to reduce the incidence of IMD, effective polysaccharide-conjugate vaccines against serogroups A, C, W and Y have been developed and introduced in national vaccination programmes. These vaccines are, however, expensive and not affordable for low-income countries. Therefore, a monovalent conjugate serogroup A vaccine was produced and successfully introduced in large-scale vaccination campaigns in countries of the so-called “meningitis belt” of sub-Saharan Africa (Diomandé et al., 2015; Trotter et al., 2017), a region stretching from the Gambia and Senegal to Ethiopia (Molesworth et al., 2002).

Prior to the vaccination campaigns that started in 2010, the overall incidence of meningococcal meningitis in the region was substantially higher than anywhere else in the world, and included epidemics that occurred in the winter months between every five and twelve years (Trotter and Greenwood, 2007). Though the vaccine has been very effective at controlling meningitis epidemics caused by serogroup A, the main cause of IMD in the meningitis belt (Diomandé et al., 2015; Trotter et al., 2017), other serogroups (C, W, and X) have emerged or expanded in the region, reducing the initial impact of the vaccine. There is also concern that virulent strains circulating in the population might switch capsule or that less virulent strains not covered by the current vaccine might acquire virulence genes (Bårnes et al., 2017; Brynildsrud et al., 2019).

Both of these potential scenarios are driven by the ability of bacteria from the genus *Neisseria* to be naturally transformable and to readily recombine their DNA with one another (Obergfell and Seifert, 2015). Various mechanisms of recombination have been described in *N. meningitidis* (Joseph et al., 2011; Marri et al., 2010; Schoen et al., 2009), involving abundant and diverse repetitive DNA sequences in its chromosome. The evolutionary and epidemiological effects of recombination in *N. meningitidis* have been studied in some detail (Joseph et al., 2011; Marri et al., 2010; Retchless et al., 2018), but less work has been undertaken to describe how the extent of recombination varies both between different lineages of *N. meningitidis* and between different regions of its complete genome in a single circulating carriage population. In particular, there is little understanding of how this recombination affects the process of natural selection. This is particularly relevant in bacteria amid the disruption of population structure caused by large-scale vaccine introduction (Potts et al., 2018).

Burkina Faso, located in the center of the meningitis belt, historically has had a high burden of disease caused by serogroup A meningococci (Nicolas et al., 2005), and was one of the first countries to introduce the serogroup A conjugate vaccine in a mass vaccination campaign in 2010 (Kristiansen et al., 2013). Since then, the incidence of IMD has decreased overall, but there have been meningitis outbreaks caused by serogroups W and X, belonging, respectively, to the sequence types (STs) 11 and 181 (Kristiansen et al., 2013).

Here we present a detailed population genetic analysis, focusing on recombination, in a collection of 2848 *N. meningitidis* carriage isolates collected from three areas of Burkina Faso over the course of the implementation of the serogroup A vaccine, from 2009-2012. Whole-genome sequencing with Illumina short-read technology was used to generate *de novo* assemblies for each isolate, which were then used to construct phylogenies, infer recombination events, and perform tests for selection. Our results demonstrate that different lineages recombine at different rates, independently from differences in their population size or any possible biases in their sampling frequency. Specific recombination ‘hotspot’ regions of the genome, which undergo substantially more recombination than the rest of the genome, were found across seven of nine lineages. Some of these hotspots were shared across lineages while others were lineage-specific. This pattern of hotspots likely reflects both a non-random pattern of recombination across the *N. meningitidis* genome, and also recent routine evolution, driven by selection from the immune systems of human hosts. Recombination between lineages is primarily determined by the population size of each lineage, though the lineage’s inherent recombination rate phenotype also has an effect, particularly on the amount of incoming recombination. Finally, recombination seems to primarily be acting to reduce the effect of deleterious mutations in the genome in this population, though we also find one example of it acting to speed adaptation, potentially as a response to population expansion due to strain replacement after the vaccine. These results indicate the importance of recombination in *N. meningitidis* evolution, even in the short term, such as during the immediate aftermath of population perturbation by vaccination.

## Results

### Population structure

The PopPunk clustering of the successfully sequenced Burkina Faso collection of 2839 carriage isolates returned 17 clusters, five of which were single isolate and three had fewer than 10 isolates. These 8 clusters were not considered in the downstream per-cluster analyses, the remaining 9 clusters accounting for 99% of the isolate collection. These clusters broadly correspond to the dominant serogroups and sequence types (STs), as indicated in Figure 1; with cluster 1 being composed entirely of serogroup X and 96% of ST-181, cluster 2 99.8% of serogroup Y and 98% of ST-4375, cluster 3 composed entirely of serogroup W and 98% of ST-11, cluster 4 composed 99% of serogroup W and 93% of ST-2881, cluster 5 composed 99% of serogroup Y and 59% of ST-767, cluster 6 composed entirely of serogroup A and 99% of ST-2859, cluster 7 composed wholly of non-groupables (NG) or capsule-null (cnl) and 82% of ST-192, cluster 8 composed entirely of NG/cnl and 100% ST-198, and cluster 9 composed 100% of NG/cnl and 52% of ST-4899. Among these clusters, we found the major disease-causing lineages occurring in Burkina Faso pre (clusters 3 and 6) and post-vaccine introduction (clusters 1 and 3) (Kristiansen et al., 2013). Based on the dominant serogroups and serogroup predictions for each cluster, we can see further that the clusters contain between 1% and 24% isolates which have likely varied their capsular phenotype. Figure 2 includes the entire Burkina Faso collection, as well as the global whole-genome sequences collected as described in the methods. From it, we can see that the Burkina Faso population is composed of nine independently evolving lineages, where isolates from each cluster are more closely related to globally sampled isolates from their respective clusters as opposed to isolates sampled from other clusters within Burkina Faso.

**Fig. 1.**
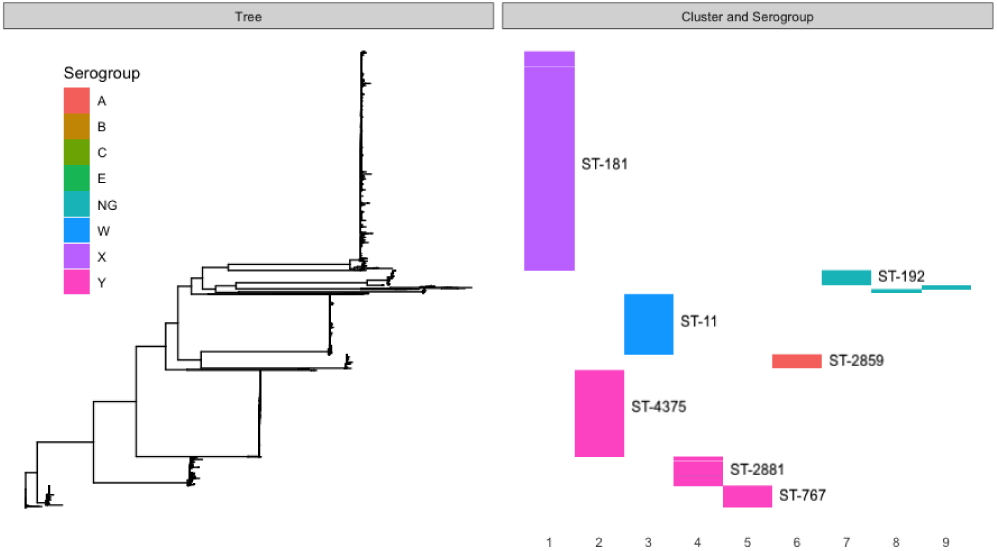
Core genome phylogeny, annotated with PopPunk-inferred clusters, serogroups, and sequence types for all Burkina Faso isolates in the collection. Clusters are numbered 1-9 from left to right, serogroups are indicated with colours as per the legend, and sequence types for the 7 largest clusters are indicated with labels. NG indicates that the isolates were non-serogroupable.

**Fig. 2.**
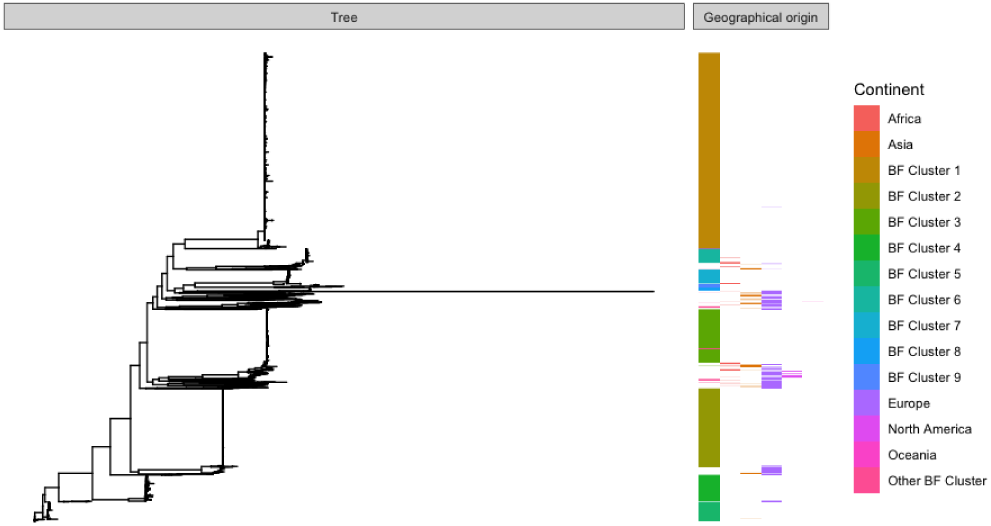
Core genome phylogeny of the Burkina Faso collection with the global background collection, arranged in order by first Burkina Faso and then alphabetical by continent, and annotated with colours for cluster, for Burkina Faso isolates, and continent of origin, for the global background collection.

### Cluster recombination rates

Average recombination event per mutation event (*ρ/θ*) rates for each cluster, as well as the overall average rate between all clusters, are shown in Table 1. These were calculated by averaging the per-branch *ρ/θ* rates inferred by Gubbins. In general, the rates were on the order of 10^−2^ with the average rate for the entire collection being 0.0745. Per-cluster average recombination rate ranged from 0.0082 to 0.1930, a difference of two orders of magnitude. The Kruskal-Wallis test on the per-branch per-mutation rates for each cluster returned an H-statistic of 440.977, with the associated *p*-value of 3.17 × 10^−90^, so there are at least two groups that are significantly different from one another. As the estimated average recombination rates are scaled to the different mutation rates of each cluster, this suggests that the clusters must be inherently different in terms of their recombination rate.

**Table 1.** Average *ρ/θ*, recombination events per mutation, for each cluster as determined by cluster-by-cluster gubbins analysis.

| Cluster (dominant serogroup/ST) | $\rho/\theta$ | Number of isolates |
| --- | --- | --- |
| Cluster 1 (X/ST-181) | 0.1053 | 1361 |
| Cluster 2 (Y/ST-4375) | 0.0082 | 539 |
| Cluster 3 (W/ST-11) | 0.0249 | 373 |
| Cluster 4 (W/ST-2881) | 0.1460 | 180 |
| Cluster 5 (Y/ST-767) | 0.0710 | 135 |
| Cluster 6 (A/ST-2859) | 0.0542 | 86 |
| Cluster 7 (NG/ST-192) | 0.0524 | 95 |
| Cluster 8 (NG/ST-198) | 0.0154 | 22 |
| Cluster 9 (NG/ST4899) | 0.1930 | 23 |
| Average | 0.0745 |  |

To determine which clusters were significantly different from one another, Dunn’s test was used for *post hoc* statistical testing. The full results from this set of pairwise tests are shown in Figure 3 with markers for a significant difference from another cluster on the top of the individual violin plots. In general, the rates of recombination span three orders of magnitude, from 10^−1^ to 10^−3^, with clusters 1, 4, and 9 being on the order of 10^−1^, cluster 2 being on the order of 10^−3^, and the remainder being on the order of 10^−2^. This post hoc difference testing suggested that there are essentially 3 recombination phenotypes, the highly recombinant lineages (and serogroups): 1 (X/ST-181), 4 (W/ST-2881), and 9 (NG/ST4899), the moderately recombinant clusters/serogroups: 3(W/ST-11), 5(Y/ST-767), 6(A/ST-2859), 7(NG/ST-192), and 8(NG/ST-198), and the relatively non-recombinant cluster 2 (Y/ST-4375).

**Fig. 3.**
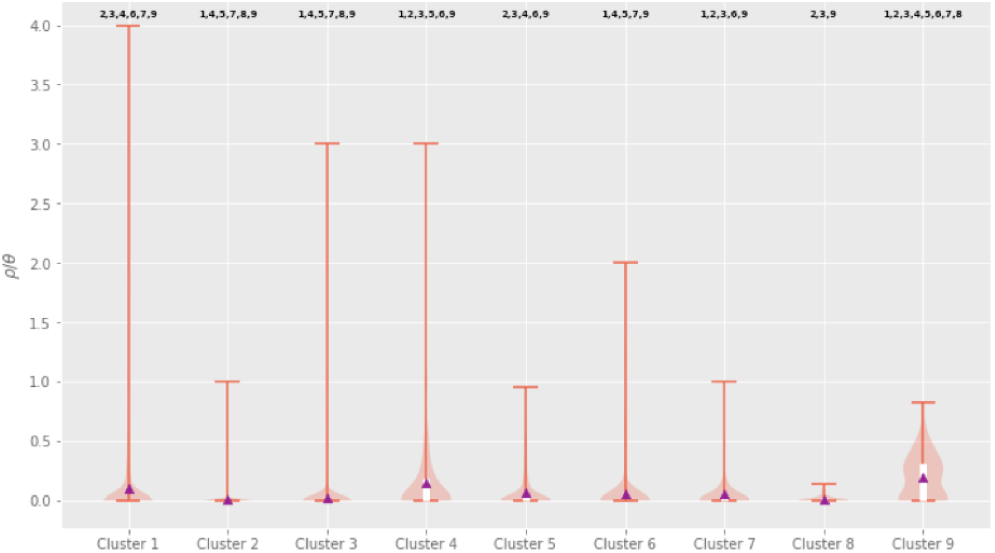
Violin plot of the per-isolate *ρ/θ*, recombination events per mutation event, as calculated by Gubbins for each cluster. The average *ρ/θ* per cluster is indicated by the purple triangles, and are also enumerated in Table 1. The top and bottom of the white boxes indicate the third and first quartiles respectively, and the whiskers of the plot represent the maximum and minimum values. The background orange background shading represents the distribution of inferred recombination rates within each cluster. Significant differences between clusters, as determined by a Kruskal-Wallis non-parametric analysis of variance on all the per-branch rates for each cluster followed by post hoc statistical testing for differences between groups using Dunn’s test and the conservative Holm-Bonferroni correction for multiple testing, are indicated by cluster numbers above each clusters’ violin plot.

### Within-Cluster Recombination hotspots

Recombination across the genome for each cluster is summarised in Figure 4, and the genes contained within each cluster’s hotspots are fully described in Table 3. From these results, we can see that in most clusters there are ‘hotspot’ regions of the genome with substantially more recombination than the background level. Seven of the clusters had obvious ‘hotspot’ peaks in the Manhattan plot of recombinations, while clusters 2 and 6 did not. Inspecting the annotation of the reference genomes for each cluster revealed a number of genes frequently present within these recombination ‘hotspots’. In particular, *pilES*, the pilin genes were present in hotspots in clusters 1, 4, 5, 7, 8, and 9. Clusters 1, 4, 5, and 7 had hotspot regions containing genes associated with iron uptake. The transferrin-binding protein gene *tbpB* was present in hotspots in clusters 1, 4, 5, and 7, and various portions of the bipartite outer membrane haemoglobin receptor gene *hpuA/B* were present in hotspots in clusters 4 and 5.

**Fig. 4.**
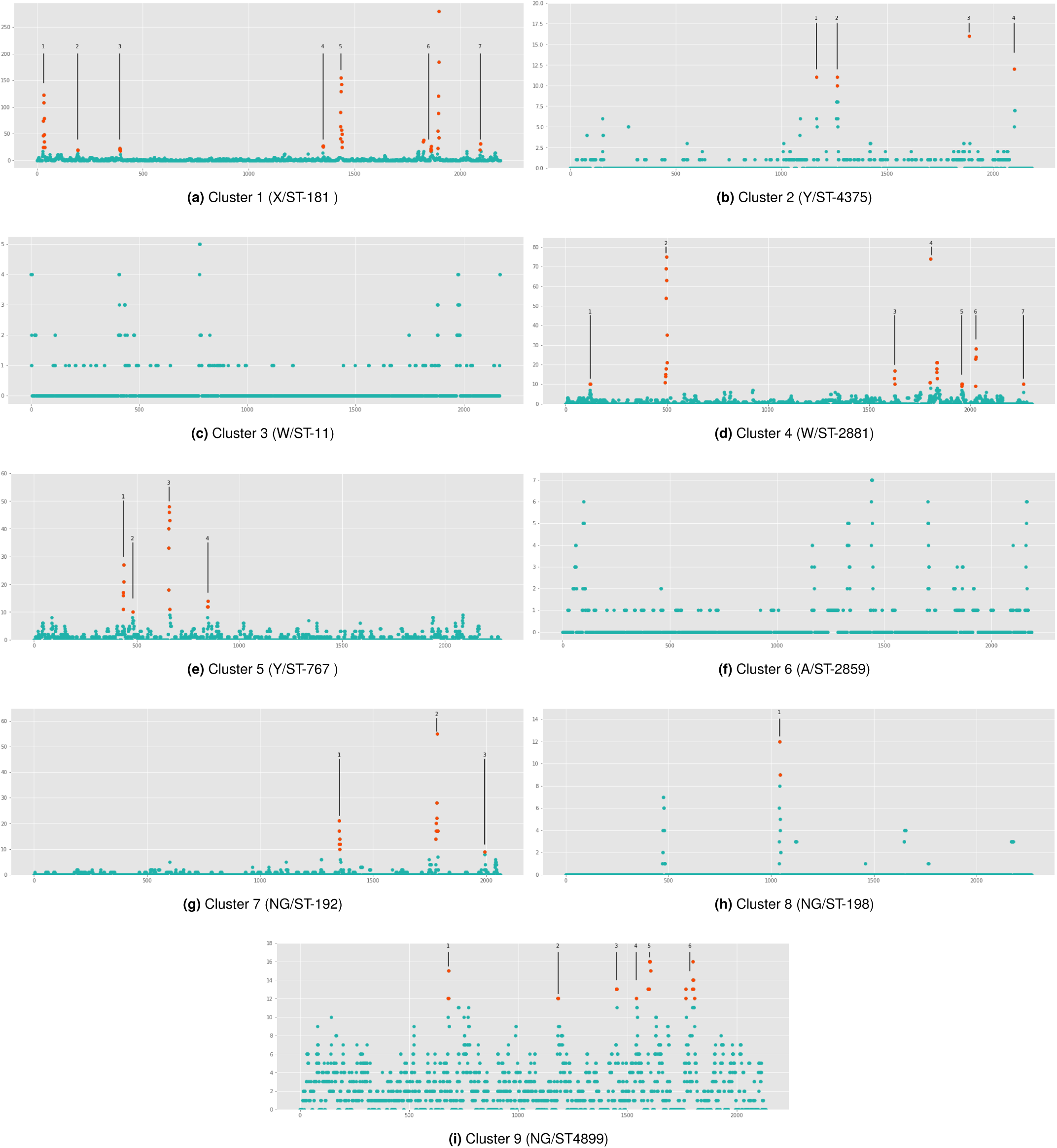
Per-cluster Manhattan plots of the number of recombinations per discrete 1000 base-pair window in each clusters’ unique reference genome. The number of recombinations in a window is indicated on the y-axis, and the position on the genome, in kilobases, is on the x-axis. Hotspots, determined manually based on obvious peaks in the plot, are highlighted in orange, and the genes within each numbered hotspot are fully described in Table 2.

### Recombination between clusters

The network of recombination events between donor and recipient clusters inferred from each gene in the pan-genome of the nine major clusters in the Burkina Faso collection is illustrated in Figure 5. A loose correlation between the size of each cluster and the number of recombinations events it was inferred to be involved in was seen, though there are exceptions to this pattern that reflect the per-cluster estimates of recombination rate for each cluster. For instance, cluster 5 was involved in more recombination events than cluster 4, despite being smaller at 135 isolates compared to 280. This is also true of the smallest clusters 8 and 9, where 9 is involved in many more recombination events than 8.

**Table 2.** Hotspots found by inspecting discrete window Manhattan plots of recombination counts for peaks. The genes contained within a single hotspot are indicated with a set of parentheses.

| Cluster (Dominant serogroup and ST) | No. of Hotspots | Genes in hotspots |
| --- | --- | --- |
| Cluster 1 (X/ST-181) | 7 | **1:** (Prolyl endopeptidase, *argA*, MJ0042 family finger-like domain gene, *pyrE*, putative uracil-DNA glycosylase, ribosomal-protein-alanine acetyltransferase, *yeaZ*, hypothetical Lipoprotein, *fda*, *xerC*, *dxs*, *miaB*) **2:** (*adhA*, *fimA*, *macA*) **3:** (*tbpB*, *lbpA*), **4:** (*glmM*, *dhpS*, *aroF*, putative AsmA-like protein) **5:** (*tbpA*, *tbp2*, *glr*, putative *yjeE*-like ATPase, *amiC*, Fic/DOC cell division protein, *rlmL*) **6:** (*ntrX*, *kinB*, *lipB*, *lipA*, *rfbC*, *rfbA*, *pilT*, *yccS*, *pilS5*, *pilE*, *pilS2*, *pilE*, *lpxC*, *yqjI*) **7:** (*porB*, *fetA*, *trmE*) |
| Cluster 2 (Y/ST-4375) | 4 | **1:** (IS1106 transposase) **2:** (*tspB*, phage-associated protein) **3:** (*lpxC*) **4:** (ISNme1 transposase, Fic/DOC hypothetical protein) |
| Cluster 3 (W/ST-11) | 0 | NA |
| Cluster 4 (W/ST-2881) | 7 | **1:** (16S rRNA, tRNA-Ile, tRNA-Ala) **2:** (Fic/DOC cell division protein, *amiC*, hypothetical ATPase, *glr*, hypothetical protein, *tbpB*, *tbpA*) **3:** (*fda*, lipoprotein, *yeaZ*, putative acetyltransferase) **4:** (*lpxC*, *pilE*, hypothetical protein) **5:** (*yccS*, *pilT*, *yggS*) **6:** (*dtpT*, hypothetical periplasmic protein, *hpuB*, *hpuA*) **7:** (*pilC*) |
| Cluster 5 (Y/ST-767) | 4 | **1:** (*pilE*, *pilS6*, *pilS5*, *pilE_2*, fimbrial protein P9-2 precursor, hypothetical protein, *pilE_3*) **2:** (*PilC*) **3:** (*tpbA*, *tpbB*, fimbrial protein P9-2 precursor), (*hpuB*, hemoglobin-haptoglobin-utilization protein) **4:** (heme oxygenase, putative lipoprotein, putative *tbp*-like) |
| Cluster 6 (A/ST-2859) | 0 | NA |
| Cluster 7 (NG/ST-192) | 3 | **1:** (*tbpA*, *tbpB*, *glr*) **2:** (*pilE_1*, *pilE_2*, *pilE_3*, *pilE_4*, *pilE_5*, *pilE_6*, hypothetical protien, *pilE_7*, *lpxC*, IS1106 transposase, IS1106A3 transposase) **3:** (hemoglobin-haptoglobin-utilization protein) |
| Cluster 8 (NG/ST-198) | 1 | **1:** (*pilE_2*, *pilE_3*, *pilE_4*, *pilS5*) |
| Cluster 9 (NG/ST-4899) | 6 | **1:** (uracil-DNA glycosylase, *pyrE*, MJ0042 family finger-like protein, *argA*) **2:** (*dapA*, hypothetical lipoprotein, pip) **3:** (*glmM*, *dhpS*, AsmA-like protein) **4:** (*aniA*, *nor*) **5:** (*lbpA*, *tbpB*, *tbpA*, *tbpB_2*) **6:** (*pilC*, *pilE_4*, *pilE_5*, *lpxC*, *yqjI*, *alx*) |

**Fig. 5.**
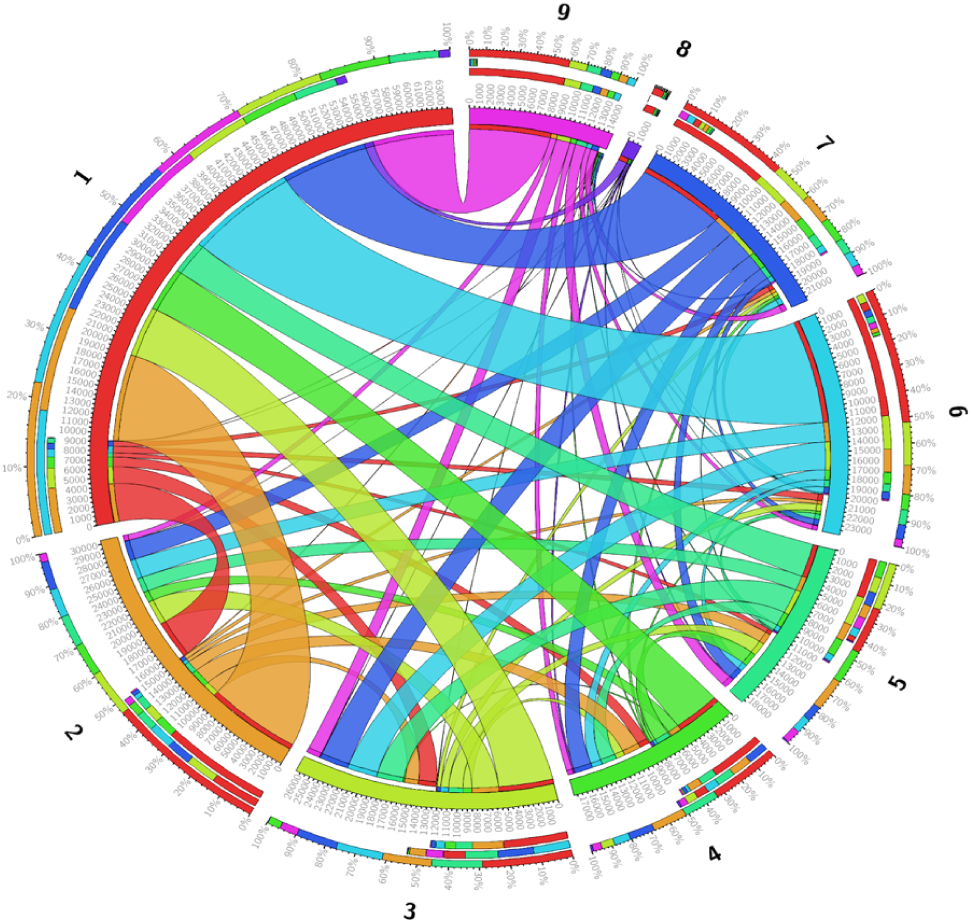
Chord diagram of the number of recombinations between the clusters in the Burkina Faso *N. meningitidis* carriage collection. Clusters are positioned on the main circle of the diagram, with the radial length of the cluster indicating the number of recombination events. Linkages between clusters represent the number of recombinations occurring between those clusters with their width representing the number of recombinations and the colour indicating the donating lineage. The three stacked bars outside the main diagram indicate, from outermost to innermost, the proportion of the total number of recombination events in each cluster coloured by the other cluster involved, those same proportions only for received recombinations, and those proportions for donated recombinations for the focal cluster.

Among the main lineages (clusters 1-9) clusters 1, 3, and 4 received more DNA by recombination than they donated, while 2, 5, 6, 7, 8, and 9 donated more DNA than they received. The results indicate that clusters with lower recombination rates and smaller clusters (eg. clusters 2, 5-9) donate more DNA in a recombination events than they receive, whereas clusters with higher recombination rate, or which are larger clusters, namely 1 and 4, and 1 and 3, receive more DNA than they donate.

Across the pan-genome of 2861 genes, the average number of recombination events per gene was 63.03, with 1084 genes not having any recombination events and the number of recombination events ranging from 1 to 9015 in genes that had inferred recombination events. It is important to remember that these ‘events’ may not directly reflect the number of actual events that occurred, particularly due to the possibility of large events being counted several times, but they should still generally reflect the amount of recombination occurring within a gene. Six genes had more than 5000 inferred recombination events, an IS200 like transposase, the flavinyl transferase *apbE*, an unnamed P1 outer membrane protein, the microcin resistance gene *tldD*, the adhesin gene *mafA*, and, the sRNA regulatory protein *yhbJ*.

### Selection and recombination in the pan-genome

*d*_*N*_/*d*_*S*_ was successfully estimated with snpgenie for 1804 genes of the 2861 genes in the pan genome, with the remaining genes being singletons or not having enough diversity to allow the calculation to be performed. The average *d*_*N*_/*d*_*S*_ for the pan-genome was 1.563, with values ranging from 0 to 396.66. Eleven genes had unusually high estimates for *d*_*N*_/*d*_*S*_ over 70 – the next highest *d*_*N*_/*d*_*S*_ estimate for a gene was 27.10 – but of these genes, six were hypothetical proteins without a known annotation, and an additional two genes were phage associated and had some pseudogenised copies. The final three genes were *hpallM* restriction enzyme, and uncharacterised peptidase, and the *exbD* membrane protein. As these genes are generally not well-described, they likely include artefacts of the annotation or pan-genome inference stage, or are simply unusual regions of the genome that do not evolve like other genes, resulting in unusually and improbably high *d*_*N*_/*d*_*S*_ estimates for these genes. Consequently, all of them were excluded from figures and further analysis.

Even with those genes excluded, correlating the *d*_*N*_/*d*_*S*_ and number of estimated recombinations in the pan-genome with a non-parametric spearman’s rank correlation returns a highly significant negative correlation coefficient of −0.122, with *p* = 2.21 × 10^−6^. As can be seen in Figure 6, however, one gene has a *d*_*N*_/*d*_*S*_ value of over 10 and 42 recombination events. This is *dpnA*, a DNA methylation modification enzyme. Additional analysis of this gene with BUSTED and FUBAR confirmed that it is under gene-wide selection (BUSTED, *p* = 0.00015), and that there is a single amino acid site under selection (FUBAR, posterior probability=0.9039). The 42 branches of the phylogeny which featured the recombination were also tested with aBSREL to see if they were under selection, and one of the branches was found to be under selection (*p* = 0.00059).

**Fig. 6.**
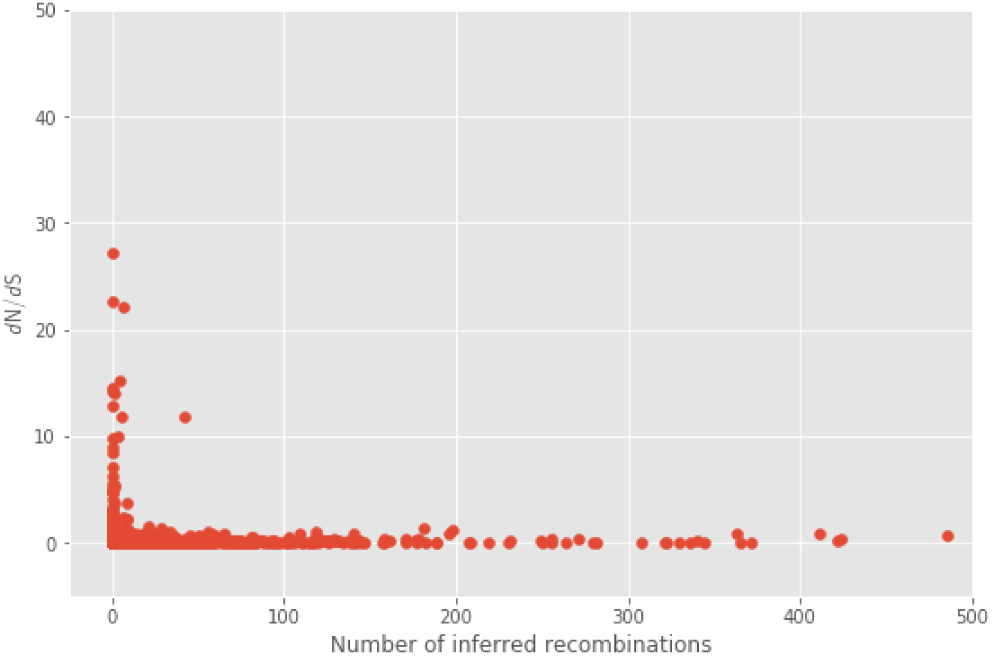
Scatter plot of number of recombination events versus *d*_*N*_/*d*_*S*_ for genes in the pan-genome of the entire Burkina Faso collection of carriage isolates. Genes with abnormally high estimates of *d*_*N*_/*d*_*S*_ were excluded from the plot as described in the text.

## Discussion

The first, immediate conclusion from the composition of our sample of *N. meningitidis* carriage isolates from Burkina Faso is that over the course of the sampling period, the population is composed primarily of nine independently evolving lineages. Bacteria of these lineages are more closely related to isolates of the same lineage from other parts of the world than to isolates from other lineages within Burkina Faso, as indicated by Figure 2, reflecting the deeply stratified population structure of *N. meningitidis*, where clonal lineages with a distant common ancestor co-exist in space and time, evolving largely independently. Seven of the lineages making up the Burkina Faso population were present in all four years of sampling. Two lineages, however, were not consistently detected. Cluster 3, the serogroup W/ST-11 lineage was first detected in our sampling in 2011, and was also detected in 2012, but not 2009 or 2010. However, it has previously been found in Burkina Faso Ouedraogo-Traore et al. (2002). Therefore, this may either represent a re-introduction of the lineage after it had gone locally extinct, or an increase in population size to be once again detectable after the population perturbation caused by vaccination. Cluster 6, the serogroup A lineage, is the other cluster which is not detected throughout our sampling period, being sampled in 2009, 2011, and 2012. That said, however, the sampled isolate in 2012 is on a long branch away from the rest of the lineage (Supp. Figure 1), and is a different sequence type, ST-7, compared to the rest of the lineage, which is ST-2859. Given that it is the target of the vaccine introduced over the course of our sampling, it seems likely that the original Serogroup A lineage in Burkina Faso went extinct by 2011 and the 2012 isolate represents a re-introduction from abroad. The dynamic changes in the composition of our population underscore the need for carriage surveillance in monitoring the evolution of *N. meningitidis* in order to guide future intervention. Though there are shifts in the population composition of our sample, including lineages which are restricted to a subset of the sampling period, a majority of lineages present are present throughout, and therefore do reflect a fluctuating yet consistent population in Burkina Faso throughout the time period of our sampling.

The significant differences in the levels of recombination between different clusters is an important and interesting result. As far as we are aware, this is the first study of a large, contemporaneous population of *N. meningitidis* that has used whole-genome data to estimate and compare the recombination rates of different lineages within the population. Though it has been shown in other species and with more disparate sampling, we show here that even within narrow geographical and temporal constraints, the lineages which make up this population of *N. meningitidis* differ in their propensity to recombine by orders of magnitude. This has important implications for how likely it is for any bacterium from a given lineage to pick up genetic material from another lineage, potentially a disease causing one. This is of particular importance with regard to the efficacy of current vaccines and the design of future ones.

The differences in within-cluster average recombination rates are additionally interesting as they suggest the rate of recombination within clusters is not tightly correlated with the clusters’ proportion of the population, and hence, stochastic opportunity, assuming that the proportions of lineages in our sample reflect those in the general population. Though recombination rate is doubtlessly affected by many factors, this does suggest that there are lineage-specific genetic components affecting the rate of recombination in *N. meningitidis*. Though there exists an understanding of the mechanisms that underlie recombination in *N. meningitidis* (Obergfell and Seifert, 2015), knowledge of how these and other genetic factors that affect the rate of recombination differ between lineages has not yet been fully explored. This awareness would not only be important to our understanding of the evolution of *N. meningitidis* and other *Neisseria*, but it would also allow for a more detailed monitoring of the populations of these serious human pathogens. Finally, it is worth noting the high rate of recombination, greatest of any cluster and more than twice the average, in one of the NG clusters in the collection, cluster 9. Non-serogroupable *N. meningitidis*, which may be entirely capsule null, or simply have loss-of-function mutations in key capsular genes, are rarely implicated in disease. However, given the extremely high recombination rate of the non-groupable cluster 9, some NG/capsule null *N. meningitidis* bacteria may play a larger role role in the evolution of a geographical population than their population frequency would suggest. Therefore, it may prove worthwhile to monitor NG/cnl *N. meningitidis* more closely in future study.

Recombination in the pan-genome of all the lineages follows a pattern that is loosely consistent with the estimated within-cluster rates. Clusters with higher estimates of average recombination rate, *ρ/θ*, are also generally involved in more within-gene recombination events in the pan genome than clusters with lower estimates of *ρ/θ*, when the population size of the cluster is taken into account. For instance, compare clusters 8 and 9, which have a similar population size but differ greatly in the number of between-cluster recombinations each is involved in. It is clear, though, that the primary factor that determines how much recombination lineages undergo in the pangenome is the proportion of each cluster within the wider population, and hence the stochastic opportunity of recombining. This is in contrast to the within-lineage recombination rates which were not correlated with population size. However, this is not unexpected given that contact between two bacteria from different clusters or lineages in a single host is less likely than contact between two bacteria of the same cluster. The recombination pattern in some clusters also suggests that the inherent recombination rate of a lineage affects how that lineage interacts with others, as cluster 2, despite being the second largest cluster, donates DNA in recombination events more often than it receives DNA in recombination events, like the much smaller clusters 5-9. This is as one would predict, as the donation of DNA in a recombination event is governed principally by the rate at which bacteria from different lineages physically encounter one another in the same host, whereas receiving DNA in a recombination event is further affected by how likely the receiving bacteria is to take up the DNA. Therefore, in lineages with low inherent recombination rates, such as cluster 2, we expect to see a bias toward donating rather than receiving of DNA, compared to what is expected for the size of the cluster. Though this effect is clearly dwarfed by the effect of cluster size, in general, this suggests that more recombinant lineages of *N. meningitidis*, at least in this population, may be able to incorporate and maintain more variation from other lineages, including from lineages that themselves are not particularly recombinant. While this may not be particularly surprising, it suggests that the inherent recombination rate has important consequences for how flexible the genome of a lineage is likely to be in response to different selection pressures, and how likely it is that it may become more virulent, antibiotic resistant, or escape a vaccine.

The radically different amounts of recombination in different regions across the genome of most of the lineages within this population, combined with the discovery of consistent genes within regions of elevated recombination in most lineages suggests two things. First, that recombination in the *N. meningitidis* genome is elevated in certain regions, and also second, that multiple lineages within the population are experiencing routine ongoing adaptation to human hosts. In particular, the elevated rates of recombination in pilin and iron uptake genes – found across clusters 1, 2, 4, 5, 7, 8, and 9 – corroborates that these genes are crucial determinants of the ability of *N. meningitidis* to survive within a human host, as pilin genes, expressed on the surface of the bacteria, are targeted by the immune system (Wachter and Hill, 2016), and human hosts are typically very iron-deficient environments for bacteria to grow in, resulting in a strong selective pressure on iron uptake pathways (Perkins-Balding et al., 2004; Yu et al., 2014). The *pilE* gene is further known to be highly recombinant, due in part to the various mechanisms of phase variation, and iron-binding proteins have been shown to be very diverse and have repeats which may promote both intra- and inter-genomic recombination close by (Acevedo et al., 2014).

The pattern of recombination events in the per-gene analysis of the pan-genome is similarly largely as expected given the within-cluster results, with an IS200 transposase, surface exposed genes like the adhesin *mafA*, and other genes that are broadly expected to be diverse, like the microcin resistance gene *tld*D in the top 5 most recombinant genes. Other genes expected to be highly recombinant, like the iron uptake genes *tbp2*/*tpbB* and the surface-exposed T-cell stimulating protein, *tspB* also have many inferred recombinations, and are within the top 15 most recombinant genes. While these genes are known to be generally very recombinant in *Neisseria*, it is also likely that the disruption of the population by vaccination has caused substantial shifts in the population size of each lineage which, in most cases apart from cluster 6, was likely a population expansion. During this demographic shift, lineages will have had more opportunity to selectively adapt to different hosts and also persist to exchange DNA, and these events are presumably contributing to the consistently high levels of recombination detected in different genes, particularly immune targets, across the different lineages.

The dN/dS values for each gene in the pan-genome follow a pattern that is broadly expected, with most values being very low, and relatively few genes – 119 – having a dN/dS > 1. It is difficult to draw specific conclusions from a simple gene-wide estimate of dN/dS, but the significant negative correlation with the number of recombinations for each gene in the pan-genome and the gene’s estimated dN/dS can inform us as to the general evolutionary role played by recombination in this population. One of the evolutionary explanations of the maintenance of recombination in populations has been that it prevents the accumulation of deleterious mutations (Felsenstein, 1974). Given the significant negative association between number of recombinations and dN/dS in the pan-genome, and the fact that non-synonymous mutations are generally more likely to be deleterious, avoiding the accumulation of deleterious mutations, or, enhancing the effect of negative selection seems to be the primary role played by recombination in this population. In spite of this, a single gene in the pan genome, the *dpnA* DNA methylation modification enzyme, has both a substantial number of recombinations, 42, and an elevated dN/dS estimate of 11.8. All of these recombinations originate from an unknown bacterium outside the collection, and 41 ended up in the cluster 1 lineage, with the final one ending up in cluster 2. Further analysis of this gene with the BUSTED algorithm for detecting gene-wide episodes of selection confirmed that it has been under selection, and using the FUBAR method for detecting specific sites under selection confirmed that the *dpnA* gene has definitely been under selection in this population. Though the specific site under selection detected by FUBAR did not overlap with the recombination, testing the branches of the gene phylogeny with the recombination present with abSREL found one of the lineages possessing the recombination to be under selection as well. This example of a high recombination gene under selection is an exception to the general pattern of recombination in the Burkina Faso population, and suggests that recombination may be acting here to speed adaptation instead of simply preventing the accumulation of deleterious mutations. Though recombination acting in this manner has long been theorised to be true for all recombining organisms under the Fisher-Muller model of recombination, and shown experimentally with plasmids in *E. coli* (Cooper, 2007), we demonstrate this phenomenon here in a sample from a natural population.

*dpnA* is also an interesting gene to have evidence of adaptation speeded by recombination in the cluster 1 lineage. Though it is uncharacterised in *N. meningitidis*, the *dpnA* gene, as part of the DpnII restriction-modification system, has been studied in *Streptococcus pneumoniae*, another bacterial pathogen that primarily colonises the nasopharynx of healthy humans. In *S. pneumoniae, DpnA* is specifically produced during competence for promoting recombination by methylating incoming DNA to protect it from the rest of the DpnII system (Johnston et al., 2013). Though we cannot assume that *dpnA* will have an identical function in *N. meningitidis*, given the important consequences DNA methylation is known to have for recombination in *N. meningitidis* (Kim et al., 2019; Seib et al., 2017) and the on-average higher rate of recombination in cluster 1 isolates with this *dpnA* recombination (0.159 vs. 0.105), it seems like that *dpnA* recombination may have affected the overall recombination phenotype in the cluster 1 lineage.

The effects of recombination on the evolution of various genes in the *N. meningitidis* genome have been the subject of extensive research. However, our understanding cannot be complete without also investigating how recombination affects the evolution of a population of *N. meningitidis* bacteria in the long term. Without this, it would be difficult to predict how different populations will respond to vaccines, anticipate changes in virulence between different lineages, or know if a population has a high risk of developing antimicrobial resistance. In this study we have conducted such an analysis on a collection of *N. meningitidis* carriage samples from Burkina Faso. We have demonstrated how using whole genome sequencing and subsequent computational analyses on whole-genome data show how it is possible to understand the differential effects of recombination on the evolution of the various lineages within this population, how the lineages of the population are interacting with each other, and the evolutionary explanations which underlie these patterns.

## Methods

### Sample collection and laboratory analysis

To assess the impact of the monovalent serogroup A conjugate vaccine on meningococcal carriage, 50,811 samples were collected over the course of four years, from 2009-2012, in 10 rounds of sampling. This took place across three sites in Burkina Faso: the Bogodogo arrondissment of the capital city, Ouagadougou; 10 villages in the Kaya district, 100km north-east of Ouagadougou, and 10 villages in the Dandé district, 350km west of Ouagadougou. Oropharyngeal swabs were taken from healthy volunteers, and associated metadata were collected from the individuals. A total of 2848 meningococcal isolates were recovered from the 10 samplings (overall carriage rate 6.05%) and confirmed as N. meningitidis at the Norwegian Institute of Public Health (NIPH), Oslo (Kristiansen et al., 2014, 2013, 2011). The isolates were serogrouped using commercial antisera (Remel, Georgia, USA), most of them were characterized by multilocus sequence typing (Maiden et al., 1998) and all were stored frozen in Greaves medium (Craven et al., 1978) at −70°C.

### Whole Genome Sequencing, quality control, assembly, and annotation

DNA was extracted from 2839 of the 2848 collected isolates and these were sent to be sequenced at the Wellcome Sanger Institute, using Illumina HiSeq 2000 sequencing technology. Paired-end libraries were prepared with insert sizes of 500 base pairs, for 125 base pair reads, sequenced at a read depth of 100x. The quality of the raw sequencing reads was assessed using an internal quality control (QC) pipeline as well as a local implementation of Kraken (Wood and Salzberg, 2014), the metagenomic sequence classifier, to check for contamination. Raw reads from 2838 isolates passed this QC and were then used to produce de novo genome assemblies using the bacterial assembly pipeline at the Sanger Institute (Page et al., 2016). The resulting genome assemblies were annotated using Prokka (Seemann, 2014), the prokaryotic annotation pipeline, to infer gene content. To identify the ST of all isolates from the sequence data, we used srst2 (Inouye et al., 2014), the Neisseria pubMLST sequence typing database, and the raw reads for each isolate. Finally, in order to determine serogroups for each isolate in silico, we adapted seroba (Epping et al., 2018) using the published capsule reference sequences (Harrison et al., 2013). In addition to the collected isolates from Burkina Faso, to give global genetic context, genomes of an additional 428 publicly-available isolates, from five continents and maximizing serogroup diversity, were downloaded from the European Nucleotide Archive and processed using the same quality control, assembly, annotation pipeline and further analyses as above.

Selected isolates from 5 of the clusters in the collection were re-sequenced with Oxford Nanopore MinION at the NIPH, with R9.4 (FLO-MIN106) flowcells and the SQK-RBK001 rapid barcoding kit for 1D reads, in order to produce references genomes for those clusters. De-multiplexing and basecalling was performed using Albacore version 2.1.2(Sahoo, 2017), and then porechop version 0.2.3(Wick et al., 2017a) was used to remove chimeric reads and adapters from the nanopore reads. The reference genomes were then assembled by hybrid assembly using both the original illumina sequence and the nanopore reads, with Unicycler (Wick et al., 2017b) before being annotated with Prokka (Seemann, 2014), like the rest of the sequenced isolates.

### Clustering and phylogeny inference

To assess the population structure of the collection of carrier isolates, we clustered the sequence assemblies using Pop-PUNK (Lees et al., 2019). For 4 clusters among the 9 largest carried forward for further analysis, an Oxford Nanopore sequenced reference was not available. Therefore, the assembly with the lowest number of contigs was selected for further scaffolding, using the MeDuSa multi-draft assembly scaf-folder (Bosi et al., 2015), and the rest of the isolate assemblies from that cluster. The resulting scaffolds for each cluster were then joined with gaps of 1000 blank nucleotides. The resulting sequences, or the Oxford Nanopre sequenced references, were then used as reference genomes in producing whole-genome alignments for each cluster. We did this using a custom mapping, variant calling, and local realignment around indels pipeline using bwa-MEM (Li, 2013), samtools mpileup (Li, 2011), and MUSCLE (Edgar, 2004), and then used the resulting whole-genome pseudo alignments to infer phylogenies for each cluster, using gubbins (Croucher et al., 2014), and RAxML, its underlying dependency (Stamatakis, 2014).

In order to construct a phylogeny for the entire collection, and also the collection combined with global isolates, we used the newly published panaroo pan-genome pipeline (Tonkin-Hill et al., 2020) to infer a set of core genes for the entire collection. We concatenated alignments of all the core genes, and then used IQ-TREE (Nguyen et al., 2014), with the substitution model, GTR+F+I+G4, inferred by ModelFinder (Kalyaanamoorthy et al., 2017), to infer phylogenies for both the entire Burkina Faso collection and the Burkina Faso collection plus global isolates.

### Recombination, selection, and pan-genome analyses

Recombination was first analysed cluster-by-cluster, using whole genome pseudo-alignments of each cluster, generated as described above, and Gubbins (Croucher et al., 2014). Gubbins outputs an estimate of *ρ/θ*, the number of recombination events per mutation event, for each branch of the phylogeny. To test if any of differences in recombination rates between clusters were statistically significant, we used a Kruskal-Wallis non-parametric analysis of variance (Kruskal and Wallis, 1952) on all of the estimated per-branch rates for each cluster, followed by Dunn’s test for post hoc statistical testing (Dunn, 1964) for differences between groups and the conservative Holm-Bonferroni correction for multiple testing (Holm, 1979). Recombination hotspot regions in the reference genome of each cluster were found by producing Manhattan-plots of the number of recombination events per 1000 base-pair discrete window across the genome, and manually looking for peaks in these plots, comparing them to a Prokka annotation of the genome to establish which genes, if any, are within these windows.

To then analyse the recombinations between clusters in the entire Burkina Faso collection, we ran fastGEAR (Mostowy et al., 2017) on alignments of each gene of the Burkina Faso collection’s pan-genome, as inferred by panaroo (Tonkin-Hill et al., 2020) and aligned using MAFFT (Katoh and Standley, 2013), using the whole-genome clustering inferred by pop-punk. fastGEAR infers recombinations explicity with directionality, allowing the flow of recombination events between clusters to be visualised as a network. This was done with Circos Table Viewer, and Circos (Krzywinski et al., 2009). To assess the effect of selection on the pan-genome, dN/dS ratios were also inferred for each gene using the same alignments as input into fastGEAR, and the implementation of the Nei-Gobujiri method (Nei and Gojobori, 1986) of calculating dN/dS as implemented in snpgenie (Nelson et al., 2015). These dN/dS results were then correlated with the number of recombination events per gene using a non-parametric spearman’s rank correlation (Spearman, 1904). Genes with an elevated snpgenie dN/dS estimate and a high number of recombinations was further analysed with the HyPhy package (Pond and Muse, 2005), in particular the BUSTED (Murrell et al., 2015) method for detecting gene-wide episodic selection, the FUBAR (Murrell et al., 2013) method for finding codons under selection, and the aBSREL (Smith et al., 2015) method to find selected branches in a phylogeny.

The scipy (Virtanen et al., 2019), pandas (McKinney et al., 2010), and matplotlib v. 2.2.3 (Hunter, 2007) python libraries were used throughout this study for statistics, data manipulation, and data visualisation, respectively. The R package ggtree (Yu et al., 2017) was used to draw phylogenies.

## DATA AVAILABILITY

All raw sequencing data generated in this study have been submitted to the European Nucleotide Archive (ENA; https://www.ebi.ac.uk/ena) under the study accession number PRJEB12668.

Metadata, assemblies, and sequence types for all the isolates in the Burkina Faso collection are available on *Neisseria* pubMLST (https://pubmlst.org/neisseria/), as per (Kristiansen et al., 2013). Whole-genome sequenced isolates are tagged with their ENA run accessions.

## ACKNOWLEDGEMENTS

Many thanks to Ingerid Kirkeleite at NIPH for DNA extraction from all the samples for Illumina and Oxford Nanopore sequencing, DNA pipelines at Wellcome Sanger Institute (WSI) for library preparation and sequencing, Pathogen informatics at WSI for systems administration support, and to the members of teams 81 and 284 at the WSI for helpful comments and advice.

## Supplementary Information

### Supplementary Figures

**Supplementary Figure 1:**
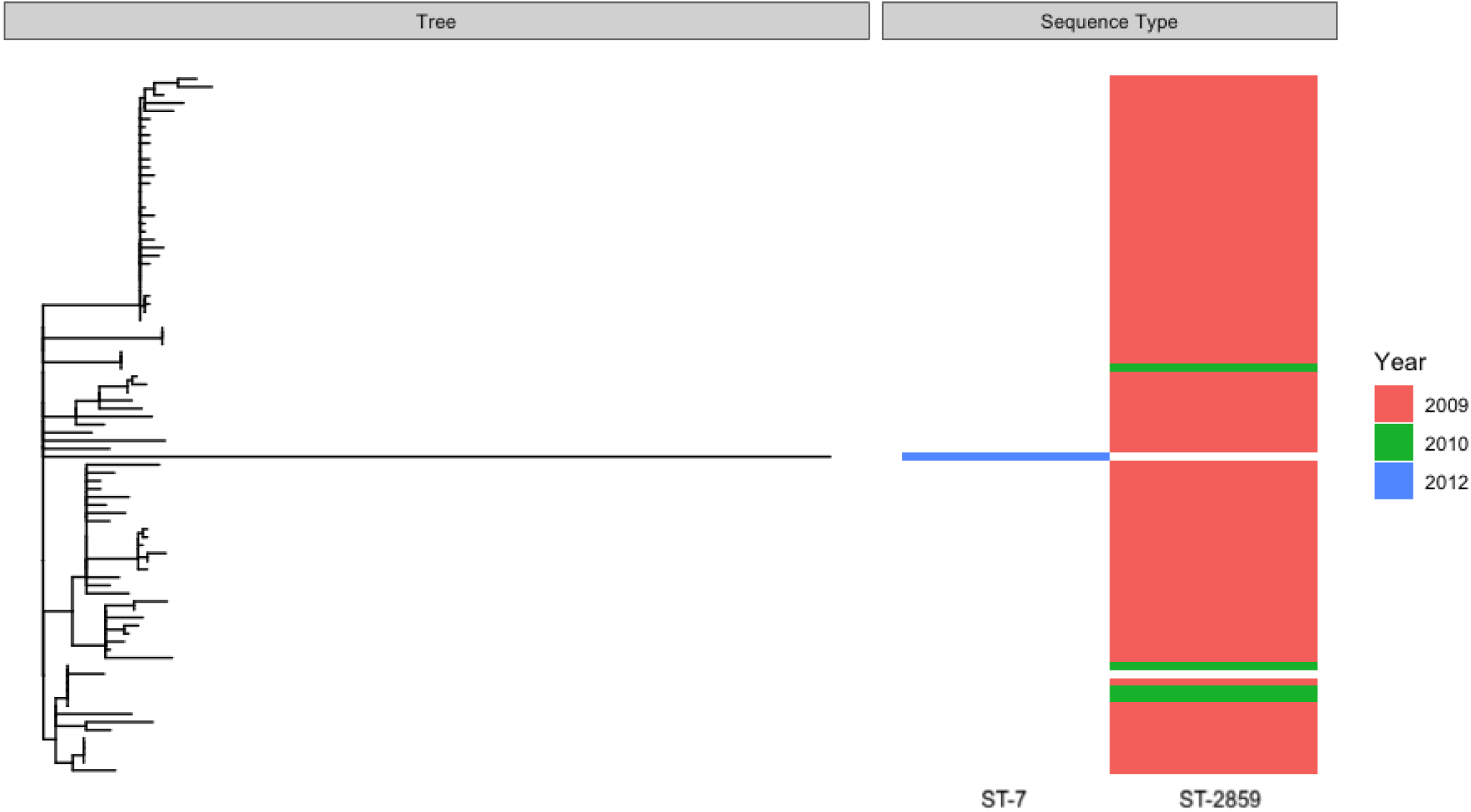
Whole genome phylogeny of the PopPUNK cluster 6, serogroup A lineage, annotated with sequence types and years of isolation. Sequence types are indicated by the column, and year of isolation by the colours as per the legend.

